# Vitamin D regulates MerTK-dependent phagocytosis in human myeloid cells

**DOI:** 10.1101/2020.02.06.937482

**Authors:** Jelani Clarke, Moein Yaqubi, Naomi C. Futhey, Sara Sedaghat, Caroline Baufeld, Manon Blain, Sergio Baranzini, Oleg Butovsky, John H. White, Jack Antel, Luke M. Healy

## Abstract

Vitamin D deficiency is a major environmental risk factor for the development of multiple sclerosis (MS). The major circulating metabolite of vitamin D (25OHD) is converted to the active form (calcitriol) by the hydroxylase enzyme *CYP27B1*. In MS lesions the tyrosine kinase MerTK expressed by microglia and macrophages regulates phagocytosis of myelin debris and apoptotic cells that can accumulate and inhibit tissue repair and remyelination. We show that calcitriol downregulates MerTK mRNA and protein expression in adult human microglia and monocyte-derived macrophages, thereby inhibiting myelin phagocytosis and apoptotic cell clearance. Proinflammatory myeloid cells express high levels of *CYP27B1* compared to homeostatic (TGFβ-treated) myeloid cells. Only proinflammatory cells in the presence of TNF-α generate calcitriol from 25OHD, resulting in repression of MerTK expression and function. The selective production of calcitriol in proinflammatory myeloid cells leading to downregulation of MerTK-mediated phagocytosis has the potential to reduce the risk for auto-antigen presentation while retaining the phagocytic ability of homeostatic myeloid cells, thereby contributing to inflammation reduction and enhanced tissue repair.

## Introduction

Vitamin D deficiency is a major environmental risk factor for the development of multiple sclerosis (MS) (1). Although widely prescribed for patients with MS, the impact of vitamin D on disease course and severity, as well as its mechanisms of action, are poorly understood. Active vitamin D (calcitriol) is obtained from the cutaneous production of vitamin D3 (cholecalciferol) in the presence of sufficient ultraviolet B irradiation, as well as limited dietary sources. Cholecalciferol is converted to 25-hydroxyvitamin D (25OHD; calcifediol), the major circulating metabolite, and then to hormonally active 1,25-dihyroxyvitamin D (1,25(OH)_2_D; calcitriol) through sequential hydroxylation, catalyzed by 25-hydroxylases (CYP2R1, CYP27A1) and 25-hydroxyvitamin D_3_ 1-alpha-hydroxylase (CYP27B1), respectively (2). Levels of 25OHD are used clinically to assess vitamin D status (3). Calcitriol functions as a ligand for the vitamin D receptor, a member of the nuclear receptor family of hormone-regulated transcription factors (3). Catabolism of 25OHD and calcitriol is initiated by the CYP24A1 enzyme, whose expression is tightly regulated by calcitriol in a negative feedback loop. CYP27B1 is abundantly expressed in most biological systems, allowing for local calcitriol production in several tissues, including the central nervous system (CNS). Importantly, CYP27B1 expression is regulated by a complex cytokine network in immune cells, including cells of myeloid origin (4).

Cells of myeloid lineage, including endogenous microglia and infiltrating monocyte-derived macrophages (MDMs), are the dominant cell population within active MS lesions (5). We have previously shown that the myeloid cell-mediated phagocytic clearance of myelin debris, a process required for efficient remyelination, is regulated by MerTK, a member of the TAM family of receptor tyrosine kinases (6). MerTK deficiency results in delayed remyelination in the cuprizone model of demyelination (7). MDMs derived from MS patients show impaired ability to phagocytose myelin, a defect linked to a reduction in MerTK expression (8). In addition to clearing myelin debris, MerTK mediates the process of efferocytosis, the removal of dead/dying cells, which is important for autoreactive T-cell fate determination in MS (9). The functions of myeloid cells are dependent on their state of activation. TGFβ, a key cytokine involved in CNS homeostasis, has been shown to maintain cells in a homeostatic state characterized by high expression of MerTK, TREM2, CSF1R and Mafb (10). In contrast, MerTK expression is comparatively lower in proinflammatory myeloid cells, a population shown to contribute to MS pathogenesis (6). Genome-wide association studies (GWAS) have explained much of MS heritability. Single nucleotide polymorphisms (SNPs) in *CYP24A1* and *CYP27B1*, which tightly regulate the intracellular levels of calcitriol have been associated with an increased risk of MS (11) (12, 13).

In the current study, we investigated calcitriol-mediated transcriptomic regulation of human MDMs and microglia. RNA sequencing revealed significant calcitriol-mediated negative regulation of both phagocytic and antigen-presenting pathways in these cell types. We demonstrate that calcitriol represses MerTK expression and phagocytic capacity of primary myeloid cells and significantly downregulates components of the antigen presentation pathway. Notably, proinflammatory myeloid cells expressing the lowest levels of MerTK have the most active vitamin D metabolic processing pathway and are therefore able to respond to the precursor 25OHD. In contrast, lack of endogenous processing of 25OHD in homeostatic myeloid cells maintains high MerTK expression and therefore participation in the immunologically-silent clearance of myelin debris and apoptotic cells.

## Methods

### Monocyte-derived macrophages

Human peripheral blood mononuclear cells (PBMCs) were isolated from healthy donors by Ficoll-Hypaque density gradient centrifugation (GE healthcare). Monocytes were isolated from PBMCs using magnetic CD14+ isolation beads (Miltenyi). Proinflammatory (MØ_GMcsf_) and alternative (M2) macrophages were generated by differentiating monocytes for 6 days in the presence of 25ng/mL GM-CSF and M-CSF respectively. To generate CNS homeostatic (MØ_0_) macrophages, TGFβ (50ng/mL) was added to the M-CSF culture conditions on days 3 and 6. 10^−7^M calcitriol (Selleckchem) was added to designated macrophages on day 1 of culture and maintained throughout differentiation. Culture media was replenished every 2-3 days.

### Microglia & astrocytes

Human adult microglia were isolated from brain tissue of patients undergoing brain surgery for intractable epilepsy. Cells were cultured in DMEM, 5% FBS, penicillin/streptomycin, and glutamine. Cell differentiation and calcitriol treatment was performed over 6 days as described above. Human fetal astrocytes were isolated as previously described (14) from human CNS tissue from fetuses at 17–23 weeks of gestation that were obtained from the University of Washington Birth defects research laboratory (BDRL, project#5R24HD000836-51) following Canadian Institutes of Health Research–approved guidelines.

### Autologous T-cells

Human T-cells were isolated from the same PBMC fraction as described for macrophages, using magnetic CD3+ isolation beads (Miltenyi Biotec).

### Proinflammatory cytokine assay

Following differentiation, macrophage cultures were supplemented with 10ng/mL TNF-α or IL-1β for 24 hours. Cells were then treated with 10^−7^ M 25OHD (Selleckchem) for 48 hours.

### Phagocytosis assay

Human myelin was isolated as previously described (24). Myelin was found to be endotoxin-free using the Limulus amebocyte lysate test (Sigma-Aldrich). To evaluate myelin uptake, myelin was incubated with a pH-sensitive dye (pHRodamine; Invitrogen) for 1h in PBS (pH 8). Dyed myelin was added to myeloid cells to a final concentration of 20ug/ml and incubated for 1h. Flow cytometry was performed using the FACS Fortessa (BD Biosciences). Live cells were gated based on live-dead staining and doublets were excluded.

### Flow cytometry

Human myeloid cells were detached gently using 2 mmol EDTA/PBS and blocked in FACS buffer supplemented with 10% normal human serum and normal mouse IgG (3 mg/ml). Cells were incubated at 4 □C for 15min with Aqua viability dye (Life Technologies) and then subsequently incubated at 4°C for 30 min with either control isotype Ab or appropriate surface marker (MerTK, CD80, CD86, HLA-DR/DP/DQ, HLA-ABC, CD40, CD274) test Abs. Cells were washed and flow cytometry was performed using the Attune NxT (Thermo Fisher Scientific). Myeloid cells were gated based on side scatter-area and forward light scatter (FSC)-area. Doublets were excluded using FSC-area and FSC-height. Live cells were gated based on live-dead staining (Aqua; Life Technologies).

### Apoptosis assay

Isolated T-cells were collected and resuspended to 1×10^6^ cells/mL in PBS. Cells were exposed to UV for 1h. Following exposure, cells were collected, pelleted, and processed for phagocytosis as previously described for myelin. pHRodamine-dyed cells were inoculated into macrophage cultures at a density of 5:1 T:M and left to incubate for 1h. Assessment of apoptosis was done by flow cytometry using Alexa 488 Annexin V/Dead cell apoptosis kit (Thermo Fisher)

### RNA sequencing

Control and calcitriol treated MDMs and microglia were collected in TRIzol reagent (Invitrogen) and RNA was extracted according to the manufacturer’s protocol (Qiagen). Smart-Seq2 libraries were prepared by the Broad Technology Labs and sequenced by the Broad Genomics Platform. cDNA libraries were generated the Smart-seq2 protocol (15). RNA sequencing was performed using Illumina NextSeq500 using a High Output v2 kit to generate 2 × 25 bp reads. Reads were aligned to the GRCh38 genome with STAR aligner and quantified by the BTL computational pipeline using Cuffquant version 2.2.1 (16, 17). Raw counts were normalized using TMM normalization and then log2-transformed. The read counts for each sample were used for differential expression analysis with the edgeR package (18, 19). The differentially-expressed genes were identified using p-value < 0.05 and log2 fold change > 1. Principle component analysis (PCA) was carried out using built-in R function, *prcomp*, and visualized using gplot package. Heatmaps were created using ggplot2 package in R. The full list of identified genes was used to generate volcano plots in R. For PCA and heatmap graphs, variance of genes across all macrophage phenotypes was calculated and the top 500 highly variable genes were used for further analysis.

### qPCR

Cells were lysed in TRIzol (Invitrogen). Total RNA extraction was performed using standard protocols followed by DNAse treatment according to the manufacturer’s instructions (Qiagen). For gene expression analysis, random hexaprimers and Moloney murine leukemia virus reverse transcriptase were used to perform standard reverse transcription. Analysis of individual gene expression was conducted using TaqMan probes to assess expression relative to *Gapdh*.

### Statistics

Paired Student’s t-test and analysis of variance, one-way ANOVA, were used to determine significance of results.

## Results

### Calcitriol mediates significant transcriptional changes in human monocyte-derived macrophages

We have previously identified MerTK as an important phagocytic receptor for the immunologically-silent clearance of myelin debris (6). To identify compounds that are known to alter *MERTK* gene expression we used a data integration approach known as iCTNet (20). iCTNet retrieves information from multiple databases and creates a single network with user-defined parameters for visualization. Calcitriol was revealed as a regulatory factor upon visualization of a sub-set of FDA-approved compounds (gray) and diseases (pink) related to *MERTK* (Fig. 1A).

**FIGURE 1.**
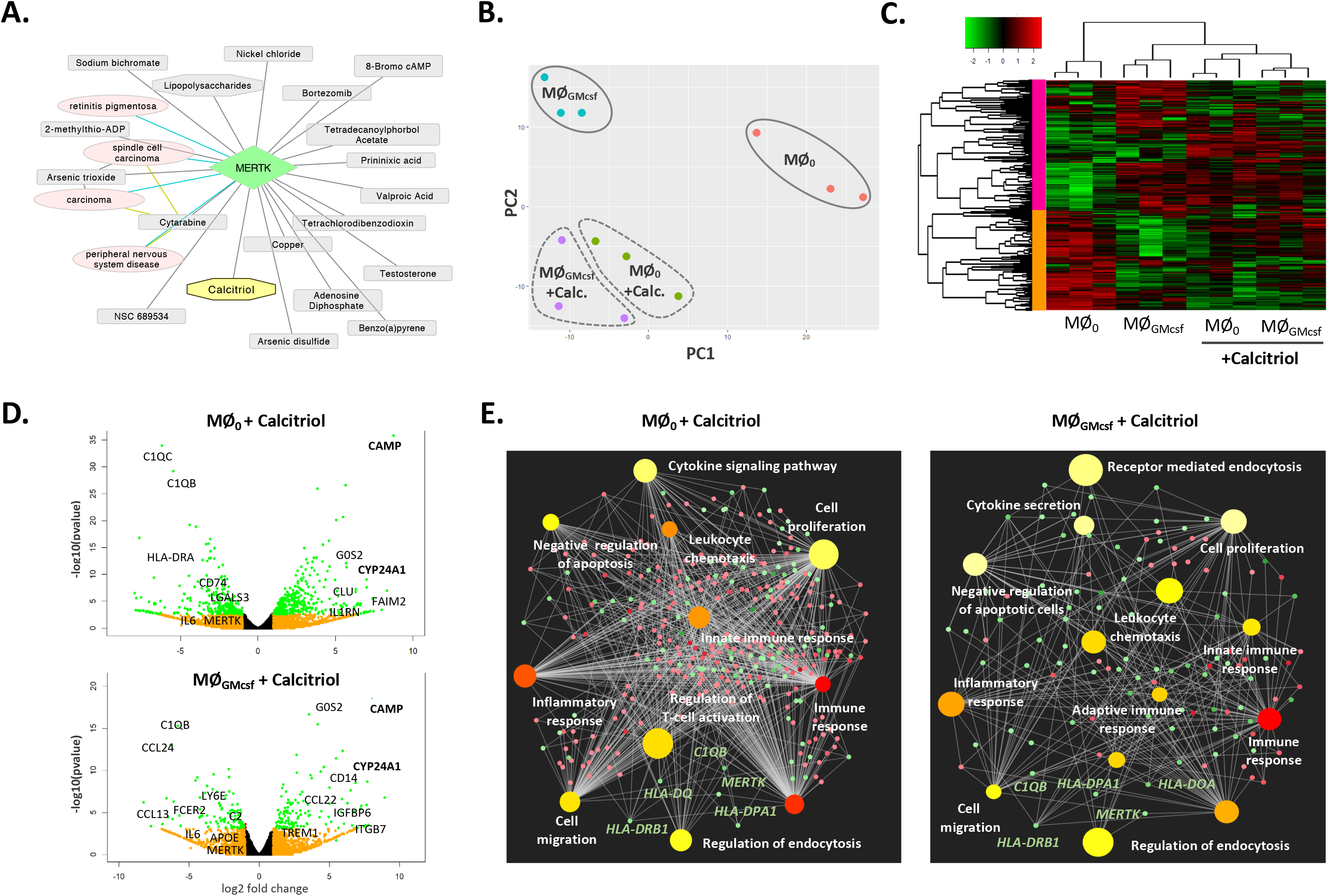
Calcitriol mediates significant transcriptional changes in human MDMs. (**A**) iCTNet neighborhood visualization of MerTK including FDA-approved compounds and diseases associated with genetic variants or mutations in MerTK. Calcitriol is identified as a MerTK-interacting molecule. (**B**) PCA plot of MØ_0_ (n3), MØ_GMcsf_ (n3), and calcitriol treated (n6) MDM samples shows separation along PC1 according to cellular phenotype and along PC2 in response to calcitriol treatment based on transcriptional profile. (**C**) Unsupervised hierarchical clustering and heat map of control and treated MDMs shows that samples cluster according to calcitriol treatment and then according to their phenotype. Upregulated genes are shown in red and downregulated genes in green. Dendrogram provides a measure of the relatedness of gene expression in each sample (top) and for each gene (left). (**D**) Volcano plots display comparison of gene expression between untreated and calcitriol treated MØ_0_ and MØ_GMcsf_ cells. Genes with adjusted p-value/FDR < 0.05 only are shown in red. Genes with log2Fold change > 1 in orange and if both requirements are met, genes appear in green. Genes of interest are marked, including genes *CYP24A1* and *CAMP,* highlighting cellular response to calcitriol. (**E**) ORA networks display the most enriched biological processes. Differentially-expressed genes (FDR < 0.05; log2Fold change > 1) in response to calcitriol were used to generate networks. Set nodes represent biological processes, which are colored based on their FDR: the most significant appears in red, set nodes with comparably higher p-value are shown in light yellow. Size of the set nodes corresponds to the number of genes associated with that biological process. Smaller nodes represent individual genes, which are colored based on their fold change (upregulation = red; downregulation = green).

To examine the effect of calcitriol on MDMs in different states of polarization (supplementary Fig. 1A and B), we analyzed the transcriptomic profile of homeostatic (MØ_0_) and proinflammatory (MØ_GMcsf_) MDMs generated *in vitro* and subjected to bulk RNA sequencing. MØ_0_ show high expression of CNS homeostatic myeloid markers such as *TREM2*, *CSF1R*, *IL10* and *Mafb* (supplementary Fig. 1C). Proinflammatory MØ_GMcsf_ cells expression signatures show typical inflammatory markers such as *IL6*, *NLRP1*, *ITGAX*, *CCL22*, *MMP9* and *ITGAX*, as well as induction of inflammatory programs involving the transcription factor *BHLHE40,* identified as part of the disease-associated transcriptomic signature (21). PCA (Fig. 1B) and heatmap (Fig. 1C) analyses showed that MØ_0_ and MØ_GMcsf_ cells cluster separately based on their phenotypes with calcitriol treated cells clustering together regardless of their starting phenotype (Fig. 1B, C). Volcano plot analysis confirms this calcitriol-mediated shift in the transcriptomic signature and highlights that both phenotypes responded to calcitriol by upregulating known calcitriol target genes *CYP24A1* and cathelicidin (CAMP) (Fig. 1D). Finally, over-representation analysis (ORA) was carried out using significantly differentially expressed genes in both MØ_0_ and MØ_GMcsf_ cells exposed to calcitriol (Fig.1E). Set nodes represent biological processes colored based on p-value (red-light yellow; most significant-least significant). The size of the node corresponds to the number of genes associated with that biological process. Smaller unlabeled nodes represent individual genes (red: upregulated; green: downregulated). Down-regulated genes of interest (*MERTK* and *HLA-DRB1*) with their link to relevant biological processes (regulation of endocytosis and adaptive immune response) are highlighted.

### Calcitriol regulation of MerTK expression and function in human MDMs

Use of the Ingenuity Pathway Analysis (IPA) bioinformatic tool highlighted ‘phagosome formation’ as one of the top canonical pathways affected by calcitriol in MDMs (supplementary Fig. 2). Visualization of this pathway highlighted the downregulation of a number of phagocytic and immune-sensing receptors including complement receptors, Fc receptors, and integrins (Fig. 2A). We identified a list of 30 genes associated with phagocytosis by myeloid cells and assessed their expression in response to calcitriol in both MØ_0_ and MØ_GMcsf_ cells (Table 1). A total of 7 genes were significantly downregulated in MØ_0_ and 4 in MØ_GMcsf_ in response to calcitriol treatment. *MERTK* was the only gene significantly downregulated in both MØ_0_ and MØ_GMcsf_ cells (Fig. 2B). We validated this RNAseq finding by RT-qPCR. Regardless of phenotype, calcitriol significantly downregulated *MERTK* mRNA (Fig. 2C) and protein expression, as measured by flow cytometry (Fig. 2D).

**FIGURE 2.**
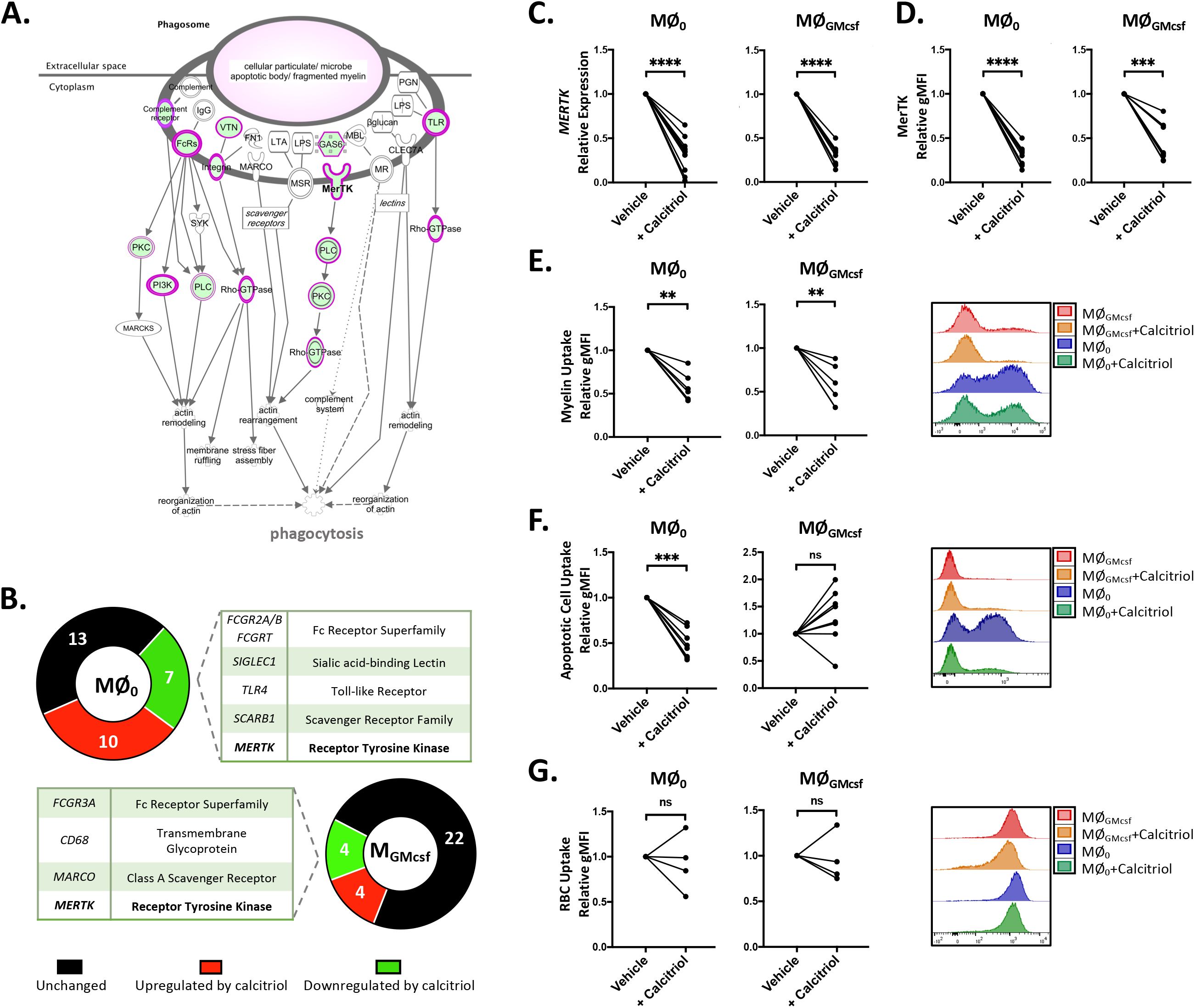
Calcitriol regulates MerTK expression and phagocytosis in human MDMs. (**A**) Ingenuity pathway analysis (IPA) of differentially-expressed genes identifies “phagosome formation” as a significantly affected pathway. Visualization of this pathway highlights affected molecules (nodes) and relationships between nodes which are denoted by lines (edges). Edges are supported by at least one reference in the Ingenuity Knowledge Base. The intensity of color in a node indicates the degree of downregulation (green). (**B**) 30 phagocytosis-related genes are identified in RNAseq datasets. Direction of regulation is assessed in both MØ_0_ and MØ_GMcsf_ cells. *MERTK* is downregulated in both cellular phenotypes. (**C**) Exposure of MDMs to calcitriol (100nM) downregulates MerTK mRNA and (**D**) protein expression in both MØ_0_ and MØ_GMcsf_ cells. (**E**) Both MØ_0_ and MØ_GMcsf_ cells are impaired in their ability to phagocytose myelin debris following treatment with calcitriol (100nM) as compared to vehicle, representative flow plot of myelin phagocytosis. (**F**) MØ_0_ cells, but not MØ_GMcsf_ cells, are impaired in their ability to phagocytose autologous apoptotic T-cells, representative flow plot of autologous apoptotic T-cell phagocytosis. (**G**) There was no significant regulation on the ability of MDMs to phagocytose opsonized red blood cells (oRBCs), representative flow plot of oRBC phagocytosis. All data was analyzed using paired Student’s t-test. *p<0.05, **p<0.01, ***p<0.001, ****p<0.0001.

To assess if reduced expression of MerTK would have a functional impact on the cells, we measured the ability of calcitriol-treated MDMs to phagocytose myelin debris, autologous apoptotic T-cells, and opsonized red blood cells (RBCs). Calcitriol-treated MDMs displayed a reduced capacity to phagocytose pHRhodamine-labelled human myelin, regardless of cellular phenotype (Fig. 2E).

In addition to myelin, MerTK has been extensively characterized as a mediator of apoptotic cell clearance (9). To investigate whether calcitriol also inhibited this process, pHRhodamine-labelled apoptotic T-cells were incubated with autologous MDMs. We observed a significant inhibition of apoptotic T-cell phagocytosis by calcitriol-exposed MØ_0_ but not MØ_GMcsf_ cells. This is indicative of a MØ_GMcsf_-specific efferocytotic receptor that can compensate for the calcitriol-mediated downregulation of MerTK. (Fig. 2F). Finally, to validate the specificity of calcitriol in regulating MerTK-dependent phagocytosis, we assessed the uptake of opsonized RBCs (oRBCs) by both MDM phenotypes. Phagocytosis of oRBCs occurs through Fc-receptor-mediated endocytosis, a MerTK-independent pathway. In all cases, calcitriol had no influence on the ability of MDMs to phagocytose oRBCs, suggesting a specificity to the calcitriol-mediated inhibition of phagocytosis by human MDMs (Fig. 2G).

### Calcitriol downregulates the expression of antigen presentation molecules

Engagement of the adaptive immune system through the re-activation of anti-myelin T-cell responses in the CNS acts as a key pathogenic step in the initiation and exacerbation of MS (22). Activation of CD8^+^ and CD4^+^ T-cells requires recognition of cognate antigens loaded on the surface of antigen-presenting cells (APCs). The strongest MS risk loci maps to the human leukocyte antigen (HLA) region, which is a gene complex encoding the major histocompatibility family of proteins (MHC). GWAS has identified the *HLA-DRB1* as the strongest risk locus, conferring a 3-fold increased MS risk (23). Activation of T-cells requires expression of MHC class molecules by APCs (signal one) in addition to a “second” signal in the form of expression of costimulatory molecules such as CD40 and CD86, both also identified as MS risk loci (24–26). IPA analysis of our sequencing results highlights the “antigen presentation pathway” as a significantly affected pathway (supplementary Fig. 2), with downregulation of both MHC class I and II molecules as indicated using the pathway visualization tool (Fig. 3A). We identified a list of 24 genes associated with antigen presentation in our dataset and assessed expression in response to calcitriol in MØ_0_ and MØ_GMcsf_ cells (Fig. 3B). Expression of a large number of *HLA/MHC* genes were downregulated by calcitriol treatment in both cellular phenotypes, including the major MS risk gene, *HLA-DRB1*. We validated these sequencing findings by measuring protein expression using flow cytometry. Protein expression of both MHC class I (HLA-ABC) and MHC class II (HLA-DR/DP/DQ) molecules were downregulated by calcitriol treatment in both MØ_0_ and MØ_GMcsf_ cells (Fig. 3C). Expression of co-stimulatory molecules CD86 and CD40 were also significantly reduced in response to calcitriol (Fig. 3D). Interestingly, we observed increased expression of immune checkpoint molecule CD274 both at the mRNA (Fig. 3B) and protein (Fig. 3E) level following treatment with calcitriol. CD274 suppresses the adaptive immune response by inducing apoptosis in CD279-expressing T-cells (27). Moreover, previous work has shown that the human *CD274* gene is a direct target of the 1,25(OH)_2_D-regulated VDR (28). Finally, a “third” signal in the form of proinflammatory cytokine release from the APC is suggested to be necessary for the induction of T-cell proliferation. IL-6 is a cytokine that when released from APCs can promote the differentiation of IL-17-producing Th-17 cells, known to be highly pathogenic in MS (29). We observed a significant decrease in IL-6 mRNA and protein release by ELISA in response to calcitriol (Fig. 3F) in MØ_GMcsf_ cells.

**FIGURE 3.**
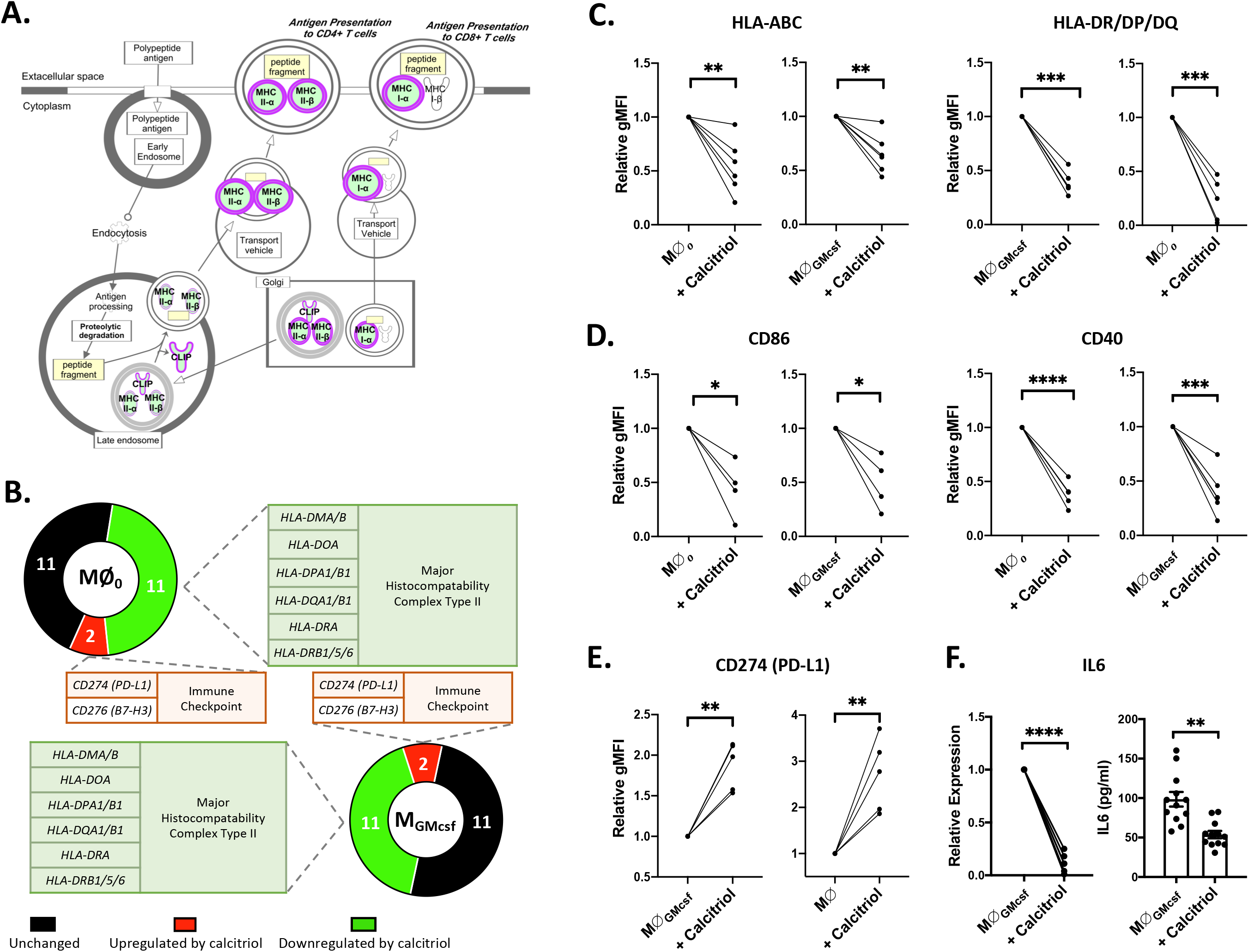
Calcitriol regulates the antigen presentation pathway in human MDMs. (**A**) Ingenuity pathway analysis (IPA) of differentially expressed genes identifies “antigen presentation” as a significantly affected pathway. Visualization of this pathway highlights affected molecules (nodes) and relationships between nodes which are denoted by lines (edges). Edges are supported by at least one reference in the Ingenuity Knowledge Base. The intensity of color in a node indicates the degree of downregulation (green). (**B**) 24 antigen presentation genes are identified in RNAseq datasets. Direction of regulation is assessed in both MØ_0_ and MØ_GMcsf_ cells with HLA genes significantly downregulated and immune checkpoint molecules upregulated in both cellular phenotypes. (**C**) Exposure of MDMs to calcitriol (100nM) downregulates protein expression of HLA-ABC and HLA-DR/DP/DQ as measured by flow cytometry (**D**) Calcitriol treatment downregulates protein expression of costimulatory molecules CD86 and CD40 in both MØ_0_ and MØ_GMcsf_ cells. (**E**) Both MØ_0_ and MØ_GMcsf_ cells upregulate CD274 (PD-L1) protein expression following treatment with calcitriol (100nM) (**F**) IL-6 mRNA and protein release (ELISA) are downregulated by calcitriol treatment in MØ_GMcsf_ cells. All data was analyzed using paired Student’s t-test. *p<0.05, **p<0.01, ***p<0.001, ****p<0.0001.

### Endogenous production of calcitriol inhibits MerTK selectively in proinflammatory MDMs

The *in vivo* circulating concentrations of calcitriol (40-100pM) are much lower than those of 25OHD (20-150nM). It is therefore important to determine whether there is sufficient intracellular metabolism of 25OHD to calcitriol within MDMs to affect MerTK expression. As shown in Fig. 4A, proinflammatory MØ_GMcsf_ cells exhibited the highest expression of the calcitriol-producing enzyme, *CYP27B1* (Fig. 4E). This high level of *CYP27B1* expression negatively correlated with *MERTK* expression. Cells that expressed the lowest levels of *MERTK* (MØ_GMcsf_) expressed the highest levels of *CYP27B1* and conversely, cells (MØ_0_) that expressed the highest levels of *MERTK* displayed the lowest expression of *CYP27B1* (Fig. 4A, B). *CYP27B1* expression is regulated by a complex network of cytokines (4); we therefore assessed the impact of proinflammatory cytokines known to play a role in MS pathology (TNF-α and IL-1β) on *CYP27B1* expression, 25OHD metabolism, and MerTK expression (30). We observed that the addition of TNF-α, and to a lesser degree IL-1β, enhanced the expression of *CYP27B1* in MØ_GMcsf_ cells but not MØ_0_ (Fig. 4C). To assess the capacity of the vitamin D metabolic pathway to regulate MerTK expression, cells were treated with the major circulating metabolite 25OHD. Despite the increased basal expression of *CYP27B1* in MØ_GMcsf_ cells, exposure to 25OHD did not significantly alter MerTK expression (Fig. 4D). However, combinatorial treatment of MDMs with 25OHD and TNF-α (and to a lesser degree IL-1β) selectively and significantly downregulated MerTK expression in MØ_GMcsf_ cells to a similar degree as calcitriol (Fig. 4D). TNF-α alone did not change MerTK expression. Altogether, we show that proinflammatory MØ_GMcsf_ cells are the only cells capable of converting 25OHD to active calcitriol, leading to the downregulation of the myelin-phagocytic receptor MerTK.

**FIGURE 4.**
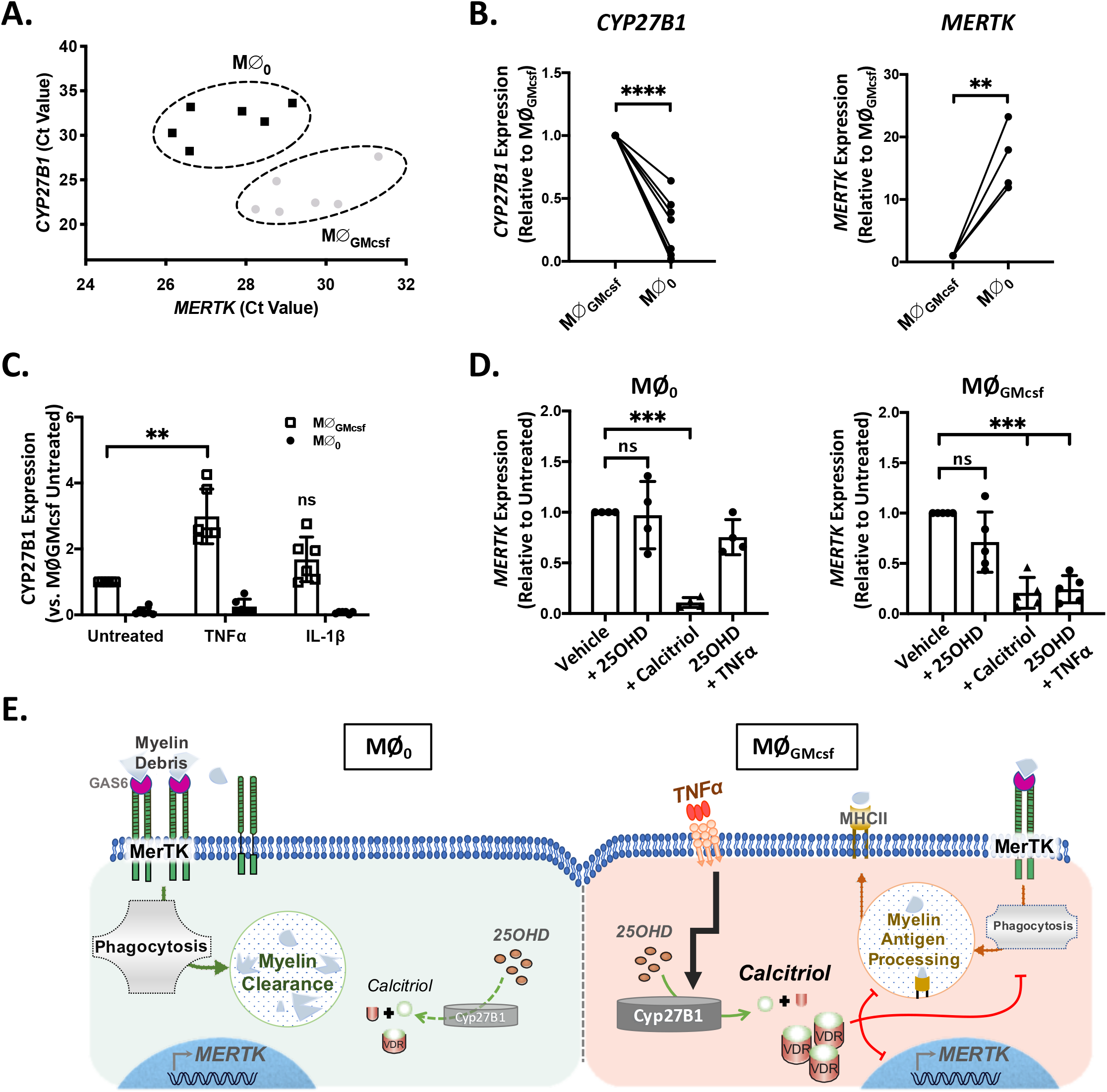
25OHD selectively downregulates MerTK in proinflammatory MDMs. (**A, B**) MØ_GMcsf_ cells express high levels of *CYP27B1* (i.e. low Ct values by qPCR) and express low levels of *MERTK*. In contrast MØ_0_ cells express the highest levels of *MERTK* and low levels of *CYP27B1*. **p<0.01, ****p<0.0001, paired Student’s t-test. (**C**) Exposure of MDMs to TNF-α and IL-1β selectively upregulates *CYP27B1* expression in MØ_GMcsf_ cells but not in MØ_0_ cells. **p<0.01, one-way ANOVA. (**D**) Combinatorial treatment of TNF-α + 25OHD selectively reduces *MERTK* expression to a similar level as calcitriol in MØ_GMcsf_ cells only. ***p<0.001, one-way ANOVA. (**E**) Schematic representation of data shows high expression of MerTK and myelin phagocytic function in homeostatic, MØ_0_ cells. These cells are unable to convert 25OHD to calcitriol due to low expression of *CYP27B1* and therefore maintain MerTK expression and function. However, proinflammatory MØ_GMcsf_ cells express high levels of *CYP27B1* and are thus able to produce calcitriol from its precursor, downregulate MerTK, molecules associated with antigen presentation, and inhibit phagocytosis.

### Calcitriol regulation of MerTK expression in primary human glia

In addition to recruited MDMs, both resident microglia and astrocyte populations take part in the neuroinflammatory process and the phagocytic clearance of myelin debris. We therefore examined the effect of calcitriol on human microglia isolated from resected brain tissue and astrocytes derived from the fetal human CNS. Microglia were polarized to CNS homeostatic (MG_0_) and proinflammatory (MG_GMcsf_) phenotypes. Similar to MDMs, cells were exposed to M-CSF (MG_0_) or GM-CSF (MG_GMcsf_) over a 6-day period with homeostatic cells receiving additional TGFβ. Confirmation of these phenotypes is highlighted by expression of established CNS homeostatic markers, including microglia-specific markers *TMEM119*, *SALL1* and *OLFML3* (supplementary Fig. 1C). Proinflammatory microglia are characterized by high expression of canonical inflammatory myeloid markers including genes that show relative specificity to microglia*, CCL17* and *IL1α* (supplementary Fig. 1D). Bulk RNA sequencing was carried out on calcitriol-treated MG_0_ and MG_GMcsf_ cells. PCA of these samples showed that, similar to MDMs, microglia cluster along the 1^st^ principal component based on their cellular phenotype (MG_0_ and MG_GMcsf_) and along the 2^nd^ principal component based on treatment with calcitriol (Fig. 5A). ORA carried out on differentially-expressed genes in both phenotypes exposed to calcitriol show a similar pattern of calcitriol-responsive biological processes, including “inflammatory response” and “cytokine production/secretion” (Fig. 5B). These transcriptomic results were validated *in vitro* whereby calcitriol downregulated *MERTK* mRNA and MerTK protein in human microglia (Fig. 5C). Finally, calcitriol had no influence on MerTK mRNA or protein expression in human fetal astrocytes (Fig. 5D), indicating that the regulation of MerTK expression by calcitriol is specific to cells of the myeloid lineage.

**FIGURE 5.**
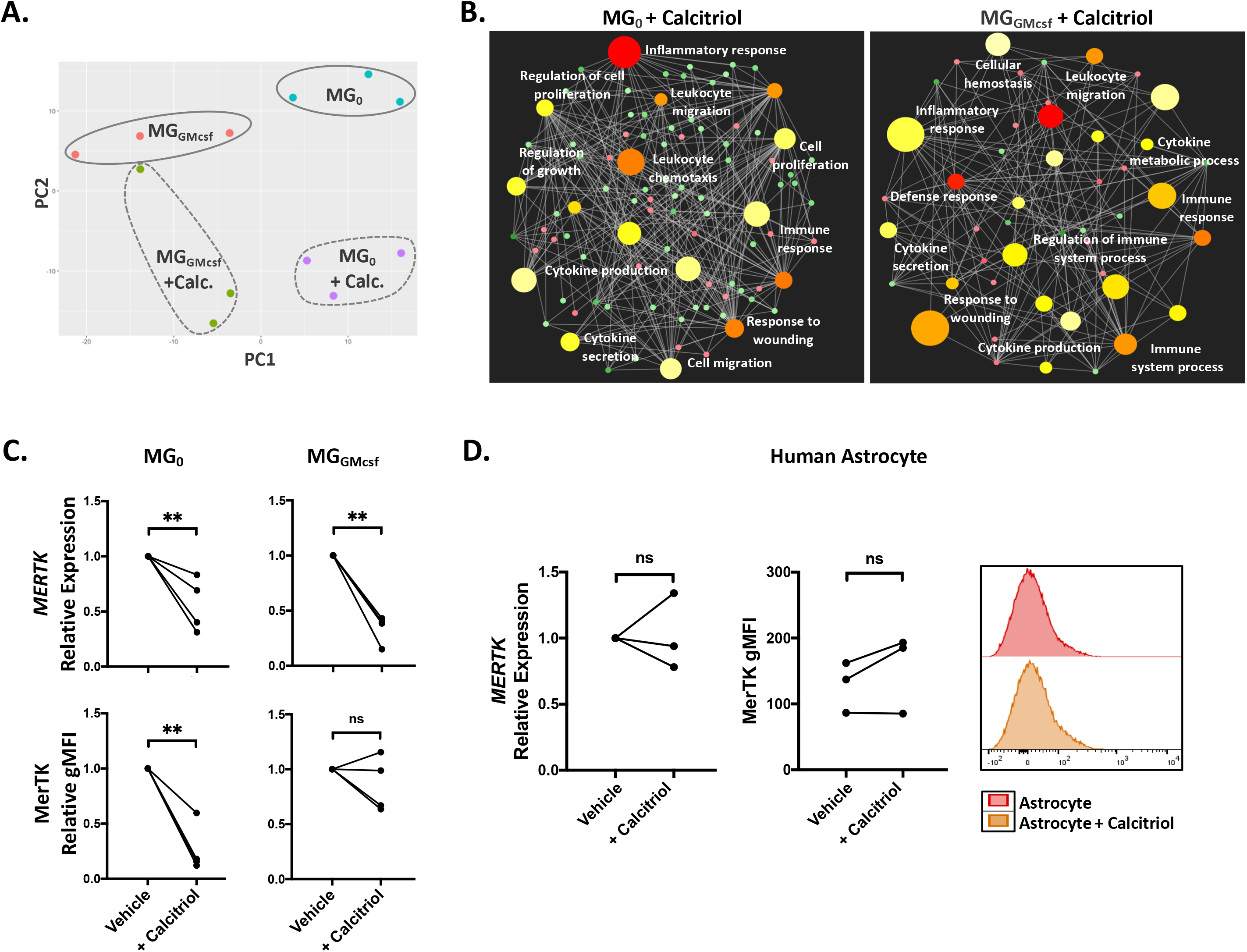
Calcitriol selectively downregulates MerTK in microglia in the brain. (**A**) Transcriptomic changes in primary human microglia (MG_0_ and MG_GMcsf_) treated with calcitriol are visualized on a PCA plot. Microglia separate along PC1 according to cellular phenotype and along PC2 in response to calcitriol treatment. (**B**) ORA networks display the most enriched biological processes. Differentially expressed genes (FDR < 0.05; log2Fold change > 1) in response to calcitriol treatment were used to generate networks. Set nodes represent biological processes, which are colored based on their FDR, the most significant appears in red, set nodes with comparably higher p-value are shown in light yellow. Size of the set nodes corresponds to the number of genes associated with that biological process. Smaller nodes represent individual genes, which are colored based on their fold change (upregulation = red; downregulation = green). (**C**) Exposure of primary human microglia to calcitriol downregulates MerTK mRNA and protein expression. **p<0.01, paired Student’s t-test (**D**) Calcitriol does not modulate MerTK mRNA or protein expression in human fetal astrocytes, representative flow plot of MerTK expression. ns = not significant, paired Student’s t-test.

## Discussion

In this study, we set out to identify compounds that could modulate expression of the phagocytic receptor tyrosine kinase MerTK, previously shown to positively regulate myelin phagocytosis and efferocytosis (6, 31). Using the cytoscape plugin iCTNet to generate a drug-gene/target network we identified calcitriol, the biologically-active form of vitamin D, as a compound that interacts with the *MERTK* gene. Here we report that calcitriol controls expression of MerTK and subsequent uptake of myelin debris and apoptotic cells by human myeloid cells. Calcitriol also establishes an immune-regulatory phenotype in these cells, significantly reducing expression of inflammatory mediators and antigen presentation machinery while increasing the expression of immune checkpoint molecules.

Myelin clearance is essential for remyelination and CNS repair (32). We and others have reported reduced MerTK expression and phagocytic capacity in myeloid cells of MS patients (33). Expression of both membrane-bound and soluble forms of MerTK are elevated in MS lesional tissues (34). In animal models, MerTK and its cognate ligand, Gas6, have shown to play protective roles, particularly in the cuprizone toxin model where Gas6-knockout mice develop a more severe level of demyelination coupled with a delayed remyelination process (35). Experimental evidence strongly supports a functional role for MerTK in inflammation resolution, debris clearance, and repair (36).

GWAS has identified several SNPs in the *MERTK* gene as independently associated with the risk of developing MS (12, 25). Fine-mapping of the *MERTK* locus identifies a risk variant that operates in *trans* with the *HLA-DRB1* locus and is associated with higher expression of MerTK in MS patient monocytes (13). This particular SNP (rs7422195) displays discordant association depending on the individual’s *HLA-DRB1*15:01* status, conferring increased risk, but converting to a protective effect on an *HLA-DRB1*15:01* homozygous background. The stratification of risk based on *DRB1* status is strongly suggestive of a functional interplay or crosstalk between phagocytosis and antigen presentation in cells capable of carrying out such functions. The beneficial role of high MerTK expression is dependent on the underlying pathology, the phase of the disease, and the activation status of the cell in which it is expressed. A recent study has shown polymorphisms in the *MERTK* gene that drive low expression of the protein in Kupffer cells to protect against the development of liver fibrosis in non-alcoholic steatohepatitis (NASH) (37).

The link between vitamin D and MS risk and the over-representation of genes involved in vitamin D metabolism as part of the genetic architecture of MS highlight the need for understanding the functional pathways under the control of vitamin D, in cell types relevant to the disease’s pathophysiology. Clinical studies are ongoing (VIDAMS & EVIDIMS), yet a reproducible benefit of vitamin D supplementation has not been evident thus far. Standard of care preparations of vitamin D are comprised of precursors such as 25OHD, the major circulating form of vitamin D. 25OHD levels define an individual’s vitamin D “status”. 25OHD is processed locally to biologically-active calcitriol (3). Four genes that control the intracellular concentration and cellular response of a cell to vitamin D are among the non-HLA MS susceptibility loci: *CYP27B1*, *CYP24A1*, *CYP2R1* and the *VDR* itself. Both systemic and intracellular conversion of 25OHD is dependent on sufficient expression of the enzyme CYP27B1 (2). Based on our findings we would predict that circulating 25OHD may not be as important as the CYP27B1-mediated production of intracellular calcitriol and subsequent transcriptional regulation of cellular function. Therefore, supplementation which increases serum 25OHD levels may not be targeting the cellular functions relevant to the pathogenesis of MS. Carlberg and Hag propose the use of a “vitamin D response index” to aid in the administration of personalized vitamin D supplementation protocols to obtain optimal vitamin D status. Such an index may assist in the future stratification of study cohorts. Based on our results we would also propose that an individual’s ability to respond to vitamin D supplementation may fluctuate with time, based on their inflammatory status and their cells’ abilities to produce active calcitriol from circulating 25OHD.

In addition to myelin debris, impaired clearance of cells undergoing apoptosis leads to sustained proinflammatory responses, as cells progress to secondary necrosis (38). Digestion of phagocytosed substrate and presentation as antigens loaded on MHC molecules (signal 1), coupled with co-stimulation (signal 2) and secretion of inflammatory cytokines (signal 3) from APCs play a critical role in stimulating the adaptive immune response (39). GWAS has identified an extended HLA haplotype, *HLA DRB1*15:01*, *DQA1*0102*, *DQB1*0602*, within the MHC class II region that is strongly associated with MS risk. In accordance with previous reports, we observed that calcitriol downregulated the expression of both MHC class I and II molecules on the surface of myeloid cells including *HLA DRB1*/*DQA1*/*DQB1*. Calcitriol downregulated the expression of major costimulatory molecules and upregulated immune checkpoint molecule CD274, as previously reported (28). Calcitriol also inhibited IL-6 expression and release. These combined data highlight the ability of calcitriol to modulate both the ingestion of material and the expression of molecular machinery involved in antigen presentation, potentially lowering the risk of auto-antigen presentation to the adaptive immune system.

Our data shows that calcitriol downregulates MerTK expression and MerTK-mediated phagocytosis in primary human myeloid cells. Intracellular production of active calcitriol from its inactive precursor and resultant repression of MerTK is limited to proinflammatory myeloid cells. This proinflammatory-specific effect may underlie a beneficial mechanism of vitamin D in MS. Proinflammatory myeloid cells are potent antigen presenters; selective inhibition of myelin uptake by these cells may lower the risk of myelin antigen presentation to infiltrating T-cells. In contrast, maintenance of MerTK and therefore phagocytic function in homeostatic myeloid populations (due to low expression of CYP27B1) would allow these cells to maintain clearance of myelin debris and contribute to the process of repair. Overall, we uncover a functional interaction between one of the strongest environmental modulators of MS risk (vitamin D) and the MerTK pathway that is selective to disease-relevant populations of primary human myeloid cells. Further understanding of the interplay between MS risk factors (both genetic and environmental) will be crucial in identifying both the correct pathways and the correct timing at which to target these pathways therapeutically.

### Study Approval

All studies were performed have been conducted according to Declaration of Helsinki principles and with approval of the Research Ethics Office at McGill University.

## References

1. Zhang, H. L., and J. Wu. 2010. Role of vitamin D in immune responses and autoimmune diseases, with emphasis on its role in multiple sclerosis. Neurosci Bull 26: 445–454.

2. Prosser, D. E., and G. Jones. 2004. Enzymes involved in the activation and inactivation of vitamin D. Trends in biochemical sciences 29: 664–673.

3. Wierzbicka, J., A. Piotrowska, and M. A. Zmijewski. 2014. The renaissance of vitamin D. Acta biochimica Polonica 61: 679–686.

4. Noyola-Martinez, N., L. Diaz, V. Zaga-Clavellina, E. Avila, A. Halhali, F. Larrea, and D. Barrera. 2014. Regulation of CYP27B1 and CYP24A1 gene expression by recombinant pro-inflammatory cytokines in cultured human trophoblasts. J Steroid Biochem Mol Biol 144 Pt A: 106–109.

5. Croxford, A. L., S. Spath, and B. Becher. 2015. GM-CSF in Neuroinflammation: Licensing Myeloid Cells for Tissue Damage. Trends in immunology 36: 651–662.

6. Healy, L. M., G. Perron, S.-Y. Won, M. A. Michell-Robinson, A. Rezk, S. K. Ludwin, C. S. Moore, J. A. Hall, A. Bar-Or, and J. P. Antel. 2016. MerTK Is a Functional Regulator of Myelin Phagocytosis by Human Myeloid Cells. The Journal of Immunology 196: 3375.

7. Binder, M. D., H. S. Cate, A. L. Prieto, D. Kemper, H. Butzkueven, M. M. Gresle, T. Cipriani, V. G. Jokubaitis, P. Carmeliet, and T. J. Kilpatrick. 2008. Gas6 Deficiency Increases Oligodendrocyte Loss and Microglial Activation in Response to Cuprizone-Induced Demyelination. The Journal of Neuroscience 28: 5195–5206.

8. Healy, L. M., J. H. Jang, S.-Y. Won, Y. H. Lin, H. Touil, S. Aljarallah, A. Bar-Or, and J. P. Antel. 2017. MerTK-mediated regulation of myelin phagocytosis by macrophages generated from patients with MS. Neurology - Neuroimmunology Neuroinflammation 4.

9. Scott, R. S., E. J. McMahon, S. M. Pop, E. A. Reap, R. Caricchio, P. L. Cohen, H. S. Earp, and G. K. Matsushima. 2001. Phagocytosis and clearance of apoptotic cells is mediated by MER. Nature 411: 207.

10. Butovsky, O., M. P. Jedrychowski, C. S. Moore, R. Cialic, A. J. Lanser, G. Gabriely, T. Koeglsperger, B. Dake, P. M. Wu, C. E. Doykan, Z. Fanek, L. Liu, Z. Chen, J. D. Rothstein, R. M. Ransohoff, S. P. Gygi, J. P. Antel, and H. L. Weiner. 2014. Identification of a unique TGF-beta-dependent molecular and functional signature in microglia. Nature neuroscience 17: 131–143.

11. Rivera, C. R., J. M. Kollman, J. K. Polka, D. A. Agard, and R. D. Mullins. 2011. Architecture and assembly of a divergent member of the ParM family of bacterial actin-like proteins. The Journal of biological chemistry 286: 14282–14290.

12. Ma, G. Z., J. Stankovich, Australia, C. New Zealand Multiple Sclerosis Genetics, T. J. Kilpatrick, M. D. Binder, and J. Field. 2011. Polymorphisms in the receptor tyrosine kinase MERTK gene are associated with multiple sclerosis susceptibility. PLoS One 6: e16964.

13. Binder, M. D., A. D. Fox, D. Merlo, L. J. Johnson, L. Giuffrida, S. E. Calvert, R. Akkermann, G. Z. Ma, Anzgene, A. A. Perera, M. M. Gresle, L. Laverick, G. Foo, M. J. Fabis-Pedrini, T. Spelman, M. A. Jordan, A. G. Baxter, S. Foote, H. Butzkueven, T. J. Kilpatrick, and J. Field. 2016. Common and Low Frequency Variants in MERTK Are Independently Associated with Multiple Sclerosis Susceptibility with Discordant Association Dependent upon HLA-DRB1*15:01 Status. PLoS Genet 12: e1005853.

14. Jack, C. S., N. Arbour, J. Manusow, V. Montgrain, M. Blain, E. McCrea, A. Shapiro, and J. P. Antel. 2005. TLR signaling tailors innate immune responses in human microglia and astrocytes. Journal of immunology (Baltimore, Md. : 1950) 175: 4320–4330.

15. Picelli, S., A. K. Bjorklund, O. R. Faridani, S. Sagasser, G. Winberg, and R. Sandberg. 2013. Smart-seq2 for sensitive full-length transcriptome profiling in single cells. Nat Methods 10: 1096–1098.

16. Dobin, A., C. A. Davis, F. Schlesinger, J. Drenkow, C. Zaleski, S. Jha, P. Batut, M. Chaisson, and T. R. Gingeras. 2013. STAR: ultrafast universal RNA-seq aligner. Bioinformatics 29: 15–21.

17. Trapnell, C., A. Roberts, L. Goff, G. Pertea, D. Kim, D. R. Kelley, H. Pimentel, S. L. Salzberg, J. L. Rinn, and L. Pachter. 2012. Differential gene and transcript expression analysis of RNA-seq experiments with TopHat and Cufflinks. Nat Protoc 7: 562–578.

18. Robinson, M. D., D. J. McCarthy, and G. K. Smyth. 2010. edgeR: a Bioconductor package for differential expression analysis of digital gene expression data. Bioinformatics 26: 139–140.

19. McCarthy, D. J., Y. Chen, and G. K. Smyth. 2012. Differential expression analysis of multifactor RNA-Seq experiments with respect to biological variation. Nucleic Acids Res 40: 4288–4297.

20. Wang, L., D. S. Himmelstein, A. Santaniello, M. Parvin, and S. E. Baranzini. 2015. iCTNet2: integrating heterogeneous biological interactions to understand complex traits. F1000Research 4: 485.

21. Krasemann, S., C. Madore, R. Cialic, C. Baufeld, N. Calcagno, R. El Fatimy, L. Beckers, E. O’Loughlin, Y. Xu, Z. Fanek, D. J. Greco, S. T. Smith, G. Tweet, Z. Humulock, T. Zrzavy, P. Conde-Sanroman, M. Gacias, Z. Weng, H. Chen, E. Tjon, F. Mazaheri, K. Hartmann, A. Madi, J. D. Ulrich, M. Glatzel, A. Worthmann, J. Heeren, B. Budnik, C. Lemere, T. Ikezu, F. L. Heppner, V. Litvak, D. M. Holtzman, H. Lassmann, H. L. Weiner, J. Ochando, C. Haass, and O. Butovsky. 2017. The TREM2-APOE Pathway Drives the Transcriptional Phenotype of Dysfunctional Microglia in Neurodegenerative Diseases. Immunity 47: 566–581 e569.

22. Lopes Pinheiro, M. A., A. Kamermans, J. J. Garcia-Vallejo, B. van Het Hof, L. Wierts, T. O’Toole, D. Boeve, M. Verstege, S. M. van der Pol, Y. van Kooyk, H. E. de Vries, and W. W. Unger. 2016. Internalization and presentation of myelin antigens by the brain endothelium guides antigen-specific T cell migration. Elife 5.

23. Baranzini, S. E., and J. R. Oksenberg. 2017. The Genetics of Multiple Sclerosis: From 0 to 200 in 50 Years. Trends Genet 33: 960–970.

24. Ji, Q., L. Castelli, and J. M. Goverman. 2013. MHC class I-restricted myelin epitopes are cross-presented by Tip-DCs that promote determinant spreading to CD8(+) T cells. Nat Immunol 14: 254–261.

25. International Multiple Sclerosis Genetics, C. e. a. 2011. Genetic risk and a primary role for cell-mediated immune mechanisms in multiple sclerosis. Nature 476: 214–219.

26. Sawcer, S., R. J. M. Franklin, and M. Ban. 2014. Multiple sclerosis genetics. The Lancet Neurology 13: 700–709.

27. Freeman, G. J., A. J. Long, Y. Iwai, K. Bourque, T. Chernova, H. Nishimura, L. J. Fitz, N. Malenkovich, T. Okazaki, M. C. Byrne, H. F. Horton, L. Fouser, L. Carter, V. Ling, M. R. Bowman, B. M. Carreno, M. Collins, C. R. Wood, and T. Honjo. 2000. Engagement of the PD-1 immunoinhibitory receptor by a novel B7 family member leads to negative regulation of lymphocyte activation. J Exp Med 192: 1027–1034.

28. Dimitrov, V., M. Bouttier, G. Boukhaled, R. Salehi-Tabar, R. G. Avramescu, B. Memari, B. Hasaj, G. L. Lukacs, C. M. Krawczyk, and J. H. White. 2017. Hormonal vitamin D up-regulates tissue-specific PD-L1 and PD-L2 surface glycoprotein expression in humans but not mice. J Biol Chem 292: 20657–20668.

29. Bettelli, E., Y. Carrier, W. Gao, T. Korn, T. B. Strom, M. Oukka, H. L. Weiner, and V. K. Kuchroo. 2006. Reciprocal developmental pathways for the generation of pathogenic effector TH17 and regulatory T cells. Nature 441: 235–238.

30. Kemanetzoglou, E., and E. Andreadou. 2017. CNS Demyelination with TNF-α Blockers. Current Neurology and Neuroscience Reports 17: 36.

31. Scott, R. S., E. J. McMahon, S. M. Pop, E. A. Reap, R. Caricchio, P. L. Cohen, H. S. Earp, and G. K. Matsushima. 2001. Phagocytosis and clearance of apoptotic cells is mediated by MER. Nature 411: 207–211.

32. Lampron, A., A. Larochelle, N. Laflamme, P. Prefontaine, M. M. Plante, M. G. Sanchez, V. W. Yong, P. K. Stys, M. E. Tremblay, and S. Rivest. 2015. Inefficient clearance of myelin debris by microglia impairs remyelinating processes. J Exp Med 212: 481–495.

33. Healy, L. M., J. H. Jang, S. Y. Won, Y. H. Lin, H. Touil, S. Aljarallah, A. Bar-Or, and J. P. Antel. 2017. MerTK-mediated regulation of myelin phagocytosis by macrophages generated from patients with MS. Neurology - Neuroimmunology Neuroinflammation 4.

34. Weinger, J. G., K. M. Omari, K. Marsden, C. S. Raine, and B. Shafit-Zagardo. 2009. Up-regulation of soluble Axl and Mer receptor tyrosine kinases negatively correlates with Gas6 in established multiple sclerosis lesions. Am J Pathol 175: 283–293.

35. Binder, M. D., H. S. Cate, A. L. Prieto, D. Kemper, H. Butzkueven, M. M. Gresle, T. Cipriani, V. G. Jokubaitis, P. Carmeliet, and T. J. Kilpatrick. 2008. Gas6 deficiency increases oligodendrocyte loss and microglial activation in response to cuprizone-induced demyelination. J Neurosci 28: 5195–5206.

36. Shafit-Zagardo, B., R. C. Gruber, and J. C. DuBois. 2018. The role of TAM family receptors and ligands in the nervous system: From development to pathobiology. Pharmacol Ther 188: 97–117.

37. Cai, B., P. Dongiovanni, K. E. Corey, X. Wang, I. O. Shmarakov, Z. Zheng, C. Kasikara, V. Davra, M. Meroni, R. T. Chung, C. V. Rothlin, R. F. Schwabe, W. S. Blaner, R. B. Birge, L. Valenti, and I. Tabas. 2019. Macrophage MerTK Promotes Liver Fibrosis in Nonalcoholic Steatohepatitis. Cell Metab.

38. Szondy, Z., Z. Sarang, B. Kiss, É. Garabuczi, and K. Köröskényi. 2017. Anti-inflammatory Mechanisms Triggered by Apoptotic Cells during Their Clearance. Frontiers in Immunology 8: 909.

39. Blander, J. M. 2008. Phagocytosis and antigen presentation: a partnership initiated by Toll-like receptors. Annals of the rheumatic diseases 67 Suppl 3: iii44–49.

