## Supplemental figures 1 & 2 for "Vitamin D regulates MerTK-dependent phagocytosis in human myeloid cells"

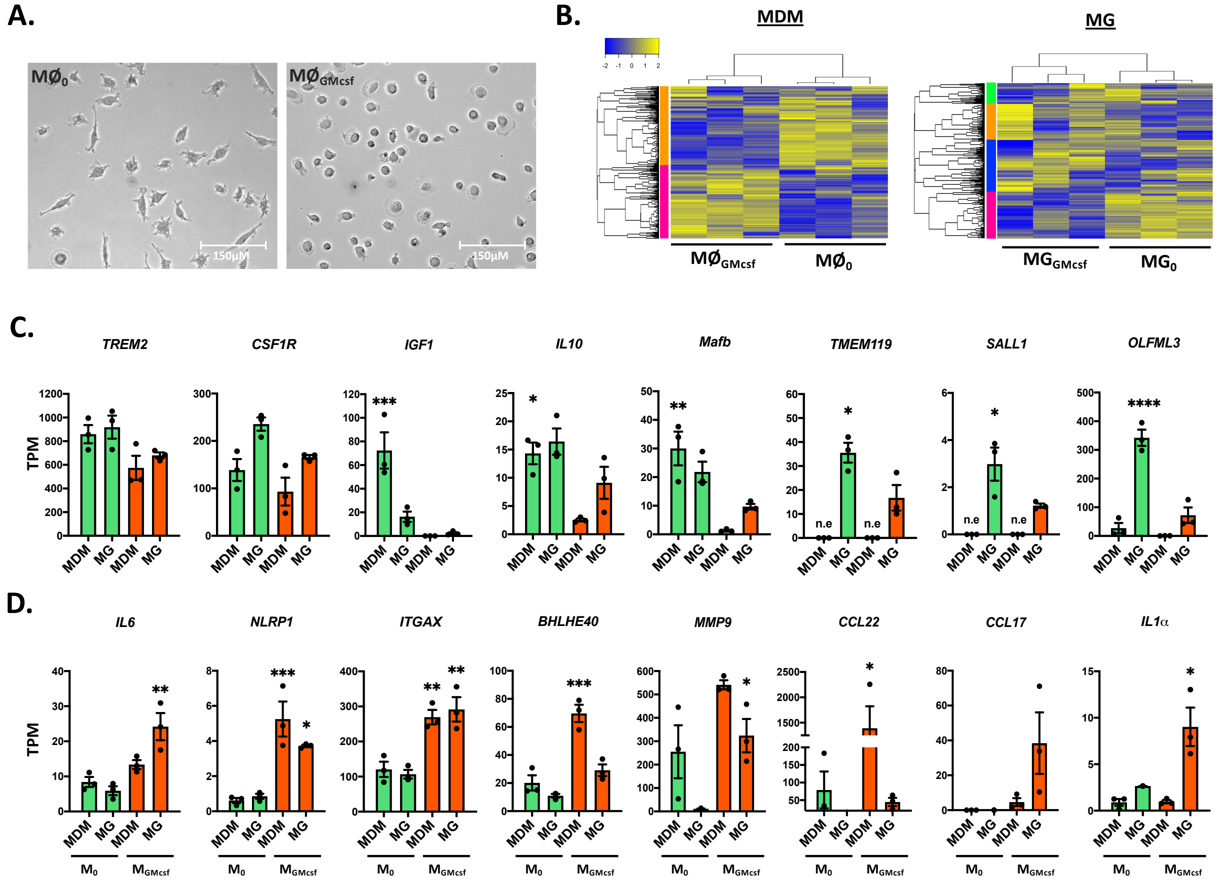


**SUPPLEMENTARY FIGURE 1.** *Confirmation of myeloid cell phenotypes*. MDMs and microglia were polarized in homeostatic (MØ$_0$ and MG$_0$) and proinflammatory (MØ$_{GMcsf}$ and MG$_{GMcsf}$) phenotypes. (**A**) shows typical morphological difference between the phenotypes. MØ$_0$ display a bipolar morphology, MØ$_{GMcsf}$ have a more ameboid and activated morphology. (**B**) Heat map of RNAseq results of polarized MDMs and microglia shows significant transcriptional differences between the two phenotypes, in both cell types. (**C**) Homeostatic myeloid cells show higher expression of known brain homeostatic markers *TREM2*, *CSF1R*, *IL10* and *Mafb*. Homeostatic microglia show increased or exclusive expression of specific homeostatic microglia markers such as *TMEM119*, *SALL1* and *OLFML3* (**D**) Proinflammatory myeloid cells show higher expression of markers of inflammation *IL6*, *NLRP1*, *ITGAX*, *BHLHE40* and *MMP9*. Some inflammatory markers are enriched in microglia populations (*CCL17* and *IL1α*) with others enriched in MDMs (*CCL22*)

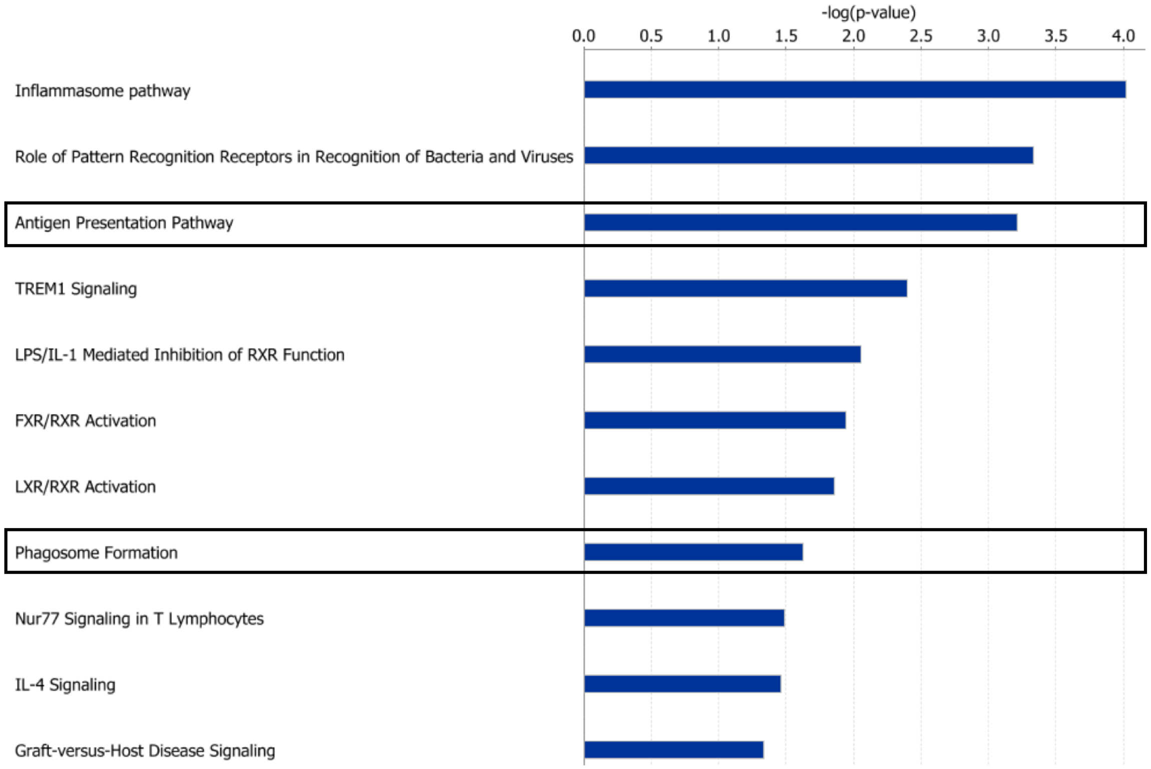


**SUPPLEMENTARY FIGURE 2.** *Pathway analysis of calcitriol-treated myeloid cells* (**A**) Pathway analysis using IPA was carried out on common significantly- and differentially-expressed genes in response to calcitriol treatment.
